# Outbreak of H9N2 avian influenza viruses in lesser rhea in Peru, June-July 2025

**DOI:** 10.64898/2026.05.08.723762

**Authors:** Alejandra Garcia-Glaessner, Alvin Crespo-Bellido, Breno Muñoz-Saavedra, Diana Juarez, Patricia Barrera, Gabriela Salmon-Mulanovich, Shadam E. Checahuari-Jaratai, Dany Cruz, Dennis Huisa, Grover Idme, Martha I. Nelson, Jesus Lescano, Mariana Leguia

**Author notes:** These authors contributed equally to this work (Alejandra Garcia-Glaessner, Alvin Crespo-Bellido).

## Abstract

Avian influenza viruses (AIVs) are endemic in the Americas and responsible for outbreaks in both domestic and wild birds that occasionally spill over into humans. We report the first known outbreak of AIV H9N2 in lesser rhea (*Rhea pennata*), also known as Darwin’s rhea, in the region of Puno-Peru. The animals in this study lived in an isolated conservation center located in remote highlands above 4,000 m.a.s.l. Between June and July 2025, a total of 46/92 animals were recorded sick, with symptoms including greenish diarrhea (100%), hyporexia (24%), dyspnea (76%), nasal discharge (42%), drowsiness (18%) and isolation from the flock (73%), and 94% later died. Gross pathology exams revealed septicemia characterized by severe hepatitis, pneumonia, tracheitis, enteritis, and encephalitis. Swab and necropsy samples tested positive for Influenza A by PCR and were later identified as H9N2 through whole genome sequencing. We generated complete H9N2 genomes for two individuals. No additional pathogens were found. Phylogenetic analysis across all eight segments revealed that the viruses were low pathogenicity H9N2 AIV strains of North American origin, which indicated this outbreak was a new introduction of the virus into South America. We also performed a comparative mutational analysis and identified multiple mutations previously associated with mammalian host adaptation, increased virulence, increased pathogenicity, and increased virus binding to α2-6 receptors, which may explain the high mortality rates observed despite the supposedly low pathogenicity of the strain. We also identified novel mutations specific to rhea viruses that will need to be experimentally validated. This is the first report of a natural H9N2 systemic infection in an avian host, highlighting a need for increased surveillance efforts for zoonotic influenza viruses with pandemic potential.

**Author Summary:** Avian influenza viruses (AIVs) are endemic in the Americas and cause more than 7,600 infections annually in domestic and wild birds worldwide each year. We report detection of AIV H9N2 in lesser rhea during an outbreak that occurred in June-July 2025 in the Andean highlands of Puno in Peru. Multiple sick animals were reported with symptoms of respiratory and gastrointestinal disease and 94% of them later died. Samples collected tested positive for Influenza A and they were subtyped as H9N2 of low pathogenic origin from North America. This is the third time H9N2 enters South America from North America, presumably through wild birds, some of which migrate along the Pacific Flyway. Comparison with other H9N2 sequences revealed a total of 44 mutations of interest that may explain the elevated death rates observed. Surveillance in wild birds remains patchy at best and needs to be strengthened in order to prevent spillover events into other animals, including humans.

## Introduction

Avian Influenza A viruses (AIV) belonging to the H9N2 subtype circulate globally in wild aquatic birds and are endemic in poultry throughout Eurasia and Africa [1]. H9N2 circulation in poultry has provided opportunities for periodic spillover into humans and other mammalian hosts, including swine, mink and cats [2–4]. The strain has also been an important donor of internal gene segments to create new reassortant genotypes with H5N1, H7N9, and H10N3 strains that have caused outbreaks in humans [5–8]. Since 2015, there have been 155 cases of AIV-H9N2 in humans, with two associated deaths [9] but no evidence of human-to-human transmission.

Low pathogenic avian influenza (LPAI) H9N2 viruses routinely circulate in North American waterfowl, but there have been no outbreaks of H9N2 in North American poultry in the last two decades. In 2005, based on serological evidence, a suspected case of H9N2 was reported in chicken in Colombia, but given a lack of virus isolation or molecular detection, confirmation of the infection remained elusive [10]. Overall, there has been very little evidence of H9N2 spillover into poultry in the Western hemisphere, especially compared to the Eastern hemisphere, where H9N2 is well adapted to domesticated birds. However, the ecology of AIV in the Western hemisphere became more complex in 2021, when highly pathogenic avian influenza (HPAI) H5N1 viruses were imported from Eurasia [11]. H5N1 rapidly spread in wild birds within North America, and shortly thereafter travelled south, reaching Latin America at the end of 2022 [12]. Starting in late 2022, Peru experienced severe HPAI-H5N1 outbreaks in seabirds and marine mammals first, and then in poultry, that resulted in mass die-offs and ecological devastation [12,13]. Between 2022 and 2025, there was frequent reassortment between H5N1 and the North American LPAI lineage [14], and in 2025 there was reassortment between H5N1 and the South American LPAI lineage that had been evolving independently for decades in waterfowl from Chile and Argentina [15]. Reassortment between North and South American lineages has also been described in wild birds [16]. Despite these events, surveillance for AIV in wildlife in Peru is almost non-existent, with mostly patchy and opportunistic sampling events occurring in response to outbreaks.

Since 1994, the Lake Titicaca Binational Special Project (PEBLT for its Spanish acronym) has promoted the conservation of natural resources in the southern Andean region of Puno, with particular focus on *Rhea pennata* (lesser rhea) (Figure 1), a large flightless bird of up to 1.7 meters in height that has been classified as critically endangered by the Peruvian government [17,18]. In response to a critical population decline of 22% from 447 to 350 animals between 2008 and 2016 [19], the PEBLT established a Rhea Conservation Center as an initiative to increase rhea numbers through breeding under semi-captive conditions with the goal of reintroducing individuals into the wild [20,21]. Rheidae are susceptible to AIV, and although rhea viruses have never been isolated or sequenced, antibodies against H5N2, H6N8, H7N1, H7N2, and H9N2 have been detected in *Rhea americana* (greater rhea) [22]. In this study we report an outbreak of LPAI-H9N2 in 46/92 semi-captive lesser rhea that resulted in 45 deaths in Southern Peru. We include complete genomic characterization, as well as phylogenetic and mutational analyses, of the H9N2 viruses identified.

**Figure 1.**
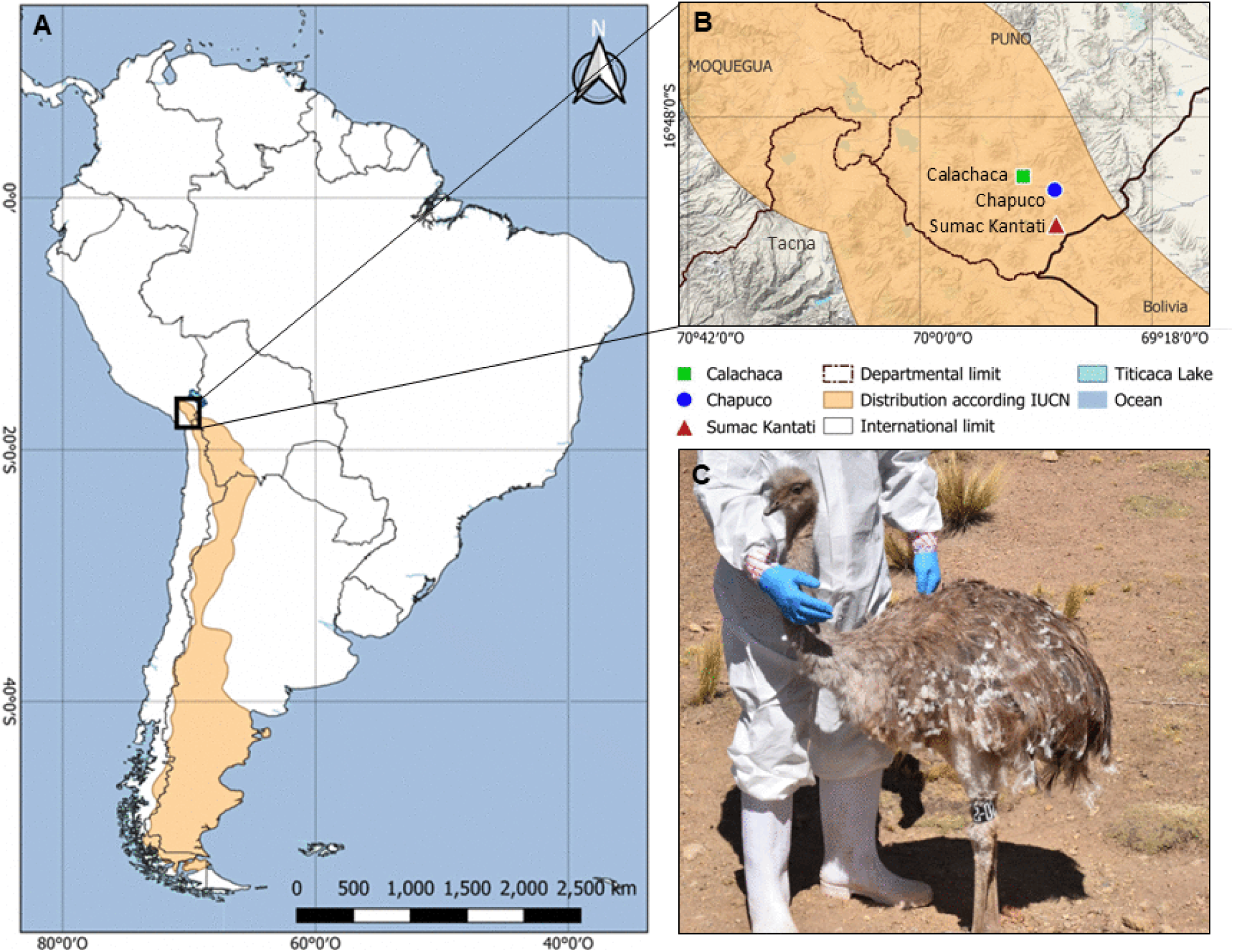
The Rhea Conservation Center in Southern Peru. (A-B) Map showing three distinct areas at different altitudes (Calachaca in green at 4,080 m.a.s.l., Chapuco in blue at 4,354 m.a.s.l. and Sumac Kantati in red at 4,401 m.a.s.l.) within the rhea conservation center in Puno where the outbreak was recorded. (C) Juevenile rhea prior to sampling in the conservation center.

## Results

### Epidemiology of the AIV H9N2 outbreak in lesser rhea

The Rhea Conservation Center, located in the Andean region of Puno, province of Collao, district of Capaso, is divided into three distinct and separate areas at different altitudes: Calachaca (4,080 m.a.s.l.), Chapuco (4,354 m.a.s.l.) and Sumac Kantati (4,401 m.a.s.l.) (Figure 1). Lesser rheas housed in two of these areas showed symptoms of digestive and respiratory disease between June and July 2025 (Figure 2). A total of 46/92 (50%) animals were recorded sick, with symptoms that included greenish diarrhea (100%), hyporexia (24%), dyspnea (76%), nasal discharge (42%), drowsiness (18%) and isolation from the flock (73%). The outbreak started June 22^nd^ in the Calachaca area, a facility that only housed juvenile animals less than two years old, where the totality of the flock (18/18) presented symptoms (100%) and 17/18 later died (94%) (Supplemental Table 1-2). By July 3^rd^, the outbreak had spread to the Chapuco area, located approximately 20 km away (∼30 km by off-road) from Calachaca, corresponding to a travel time of up to two hours depending on road conditions. This facility housed a combination of juveniles and adults, where 28/74 animals (38%) presented symptoms (6 juveniles and 22 adults) and 28/28 later died (100%). The mean survival time for juveniles and adults was 8 and 19 days, respectively, highlighting the role of age in susceptibility and suggesting that younger animals had no/very limited immunity to AIVs compared to older adults. An initial set of samples collected on July 4^th^ from four animals in Chapuco were tested by the Peruvian government for Infectious Bronchitis Virus, Avian Paramyxovirus Type 1 and AIV, and although they came back positive for Influenza A, no further testing or subtyping information became available for these samples. On July 22^nd^, additional samples were collected from two more animals in Chapuco (SER02 and SER28), including one that had to be humanely euthanized due to the severity of its symptoms, providing an opportunity to sample tissues. Gross pathology exams of the necropsied tissues revealed septicemia characterized by severe hepatitis, pneumonia, tracheitis, enteritis, and encephalitis (not shown). The samples from July 22^nd^ are the basis for this report.

**Figure 2.**
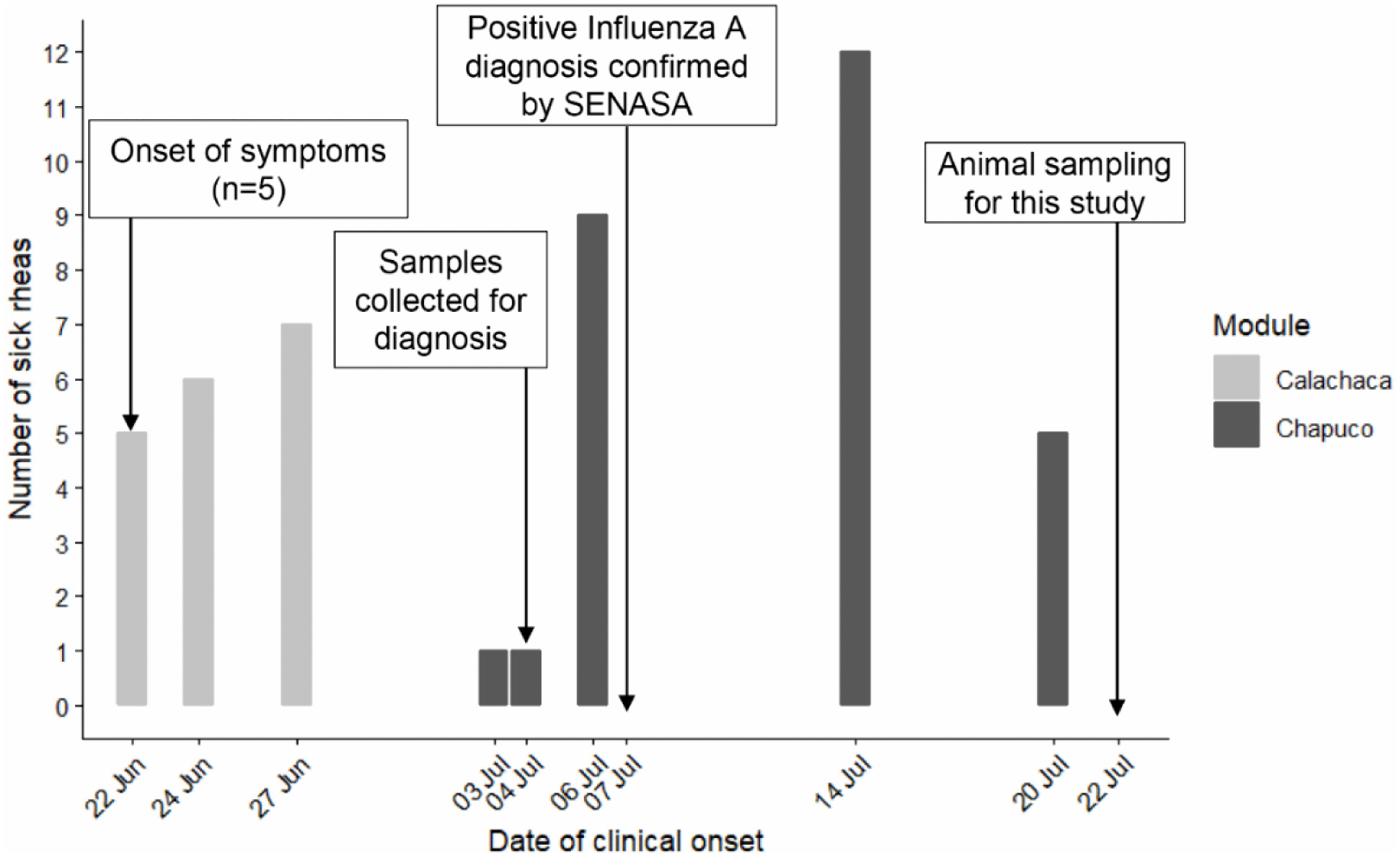
Timeline of AIV outbreak in lesser rhea, June-July, 2025 (*n*=46). Juveniles housed in Calachaca are shown in light grey, juveniles and adults housed in Chapuco are shown in dark grey.

### Detection and full genome sequencing of AIV H9N2

All samples derived from individuals SER02 and SER28 tested positive for Influenza A using a pan-Influenza A RT-qPCR assay and generated complete genomes using an NGS pipeline specific for Influenza A (Table 1). SER02 and SER28 libraries generated 2,967,354 and 1,617,407 influenza reads, respectively. This was sufficient to reconstruct the 8 genomic segments of the influenza genome in both individuals with a minimum coverage of 35X and 158X for SER02 and SER28, respectively, and to preliminarily subtype them as LPAI-H9N2 by BLAST. Sequences (A/rhea/Peru/PUN-SER02/2025(H9N2) and A/rhea/Peru/PUN-SER28/2025(H9N2)) for all genomic segments of the two viruses have been deposited in GenBank under Accession Numbers PX488774-PX488781 and PX488766-PX488773.

**Table 1.** Peruvian lesser rhea LPAI-H9N2 samples tested in this study.

| Virus Name | Species | Date | Found | Sex | Age | Sample | Ct | GenBank Accession |
| --- | --- | --- | --- | --- | --- | --- | --- | --- |
| A/rhea/Peru/PUN-SER28/2025(H9N2) | *Rhea pennata* | 2025/07/22 | Alive | n/a | Juvenile | Pharynx | 24.11 | PX488774 - PX488781 |
| A/rhea/Peru/PUN-SER02/2025(H9N2) | *Rhea pennata* | 2025/07/22 | Alive | F | Juvenile | Cloaca | 25.84 | PX488766 - PX488773 |
| A/rhea/Peru/PUN-SER02/2025(H9N2) | *Rhea pennata* | 2025/07/22 | Alive | F | Juvenile | Pharynx | 26.02 | n/a |
| A/rhea/Peru/PUN-SER02/2025(H9N2) | *Rhea pennata* | 2025/07/22 | Alive | F | Juvenile | Gizzard | 27.45 | n/a |
| A/rhea/Peru/PUN-SER02/2025(H9N2) | *Rhea pennata* | 2025/07/22 | Alive | F | Juvenile | Kidney | 27.64 | n/a |
| A/rhea/Peru/PUN-SER02/2025(H9N2) | *Rhea pennata* | 2025/07/22 | Alive | F | Juvenile | Liver | 29.52 | n/a |
| A/rhea/Peru/PUN-SER02/2025(H9N2) | *Rhea pennata* | 2025/07/22 | Alive | F | Juvenile | Trachea | 30.08 | n/a |
| A/rhea/Peru/PUN-SER02/2025(H9N2) | *Rhea pennata* | 2025/07/22 | Alive | F | Juvenile | Brain | 31.31 | n/a |
| A/rhea/Peru/PUN-SER02/2025(H9N2) | *Rhea pennata* | 2025/07/22 | Alive | F | Juvenile | Proventriculus | 32.29 | n/a |
| A/rhea/Peru/PUN-SER02/2025(H9N2) | *Rhea pennata* | 2025/07/22 | Alive | F | Juvenile | Left Lung | 33.33 | n/a |
| A/rhea/Peru/PUN-SER02/2025(H9N2) | *Rhea pennata* | 2025/07/22 | Alive | F | Juvenile | Right Lung | 34.37 | n/a |

### H9N2 rhea viruses derive from the North American LPAI lineage

To confirm virus typing and to assess phylogenetic relationships, including possible reassortment events that might explain the high mortality rates observed, we independently inferred global phylogenies for all 8 genomic segments using both background LPAI and HPAI-H5Nx clade 2.3.4.4b sequences from the Western hemisphere and Eurasia (PB2 (*n=*44,184), PB1 (*n=*43,802), PA (*n=*44,357), H9 (*n=*18,092), NP (*n=* 4,611), N2 (*n=*1,358), MP (*n=*44,567) and NS-allele A (*n=*42,238)). HPAI H5Nx clade 2.3.4.4b was used for phylogenetic context given that, since its incursion into North America in late 2021, and later into South America in 2022, it has both become widespread in the Western hemisphere and is associated with severe neurotropic infections with high mortality rates in avian hosts, including in greater rhea [23]. Phylogenetic analysis confirmed the lesser rhea viruses as LPAI strains, and further, it indicated they were of North, rather than South American origin (Figures 3 and 4). The phylogenetic trees of all six internal gene segments show lesser rhea viruses (in red) positioned within the North American LPAI lineage (in yellow), which is distinct from the South American LPAI lineage (in blue) that is routinely detected in Argentina and Chile (Figure 3). The lesser rhea viruses also cluster in a different section of the tree than HPAI-H5N1 (B3.2 genotype) viruses (in green) introduced from North into South America in 2022. The North American LPAI lineage has entered South America multiples times through independent introduction events, as evidenced by small clusters of South American viruses nested within the larger North American LPAI lineage. Nevertheless, the Peru lesser rhea viruses are not related to any of these clusters, suggesting a new, previously undetected introduction of North American LPAI into South America.

**Figure 3.**
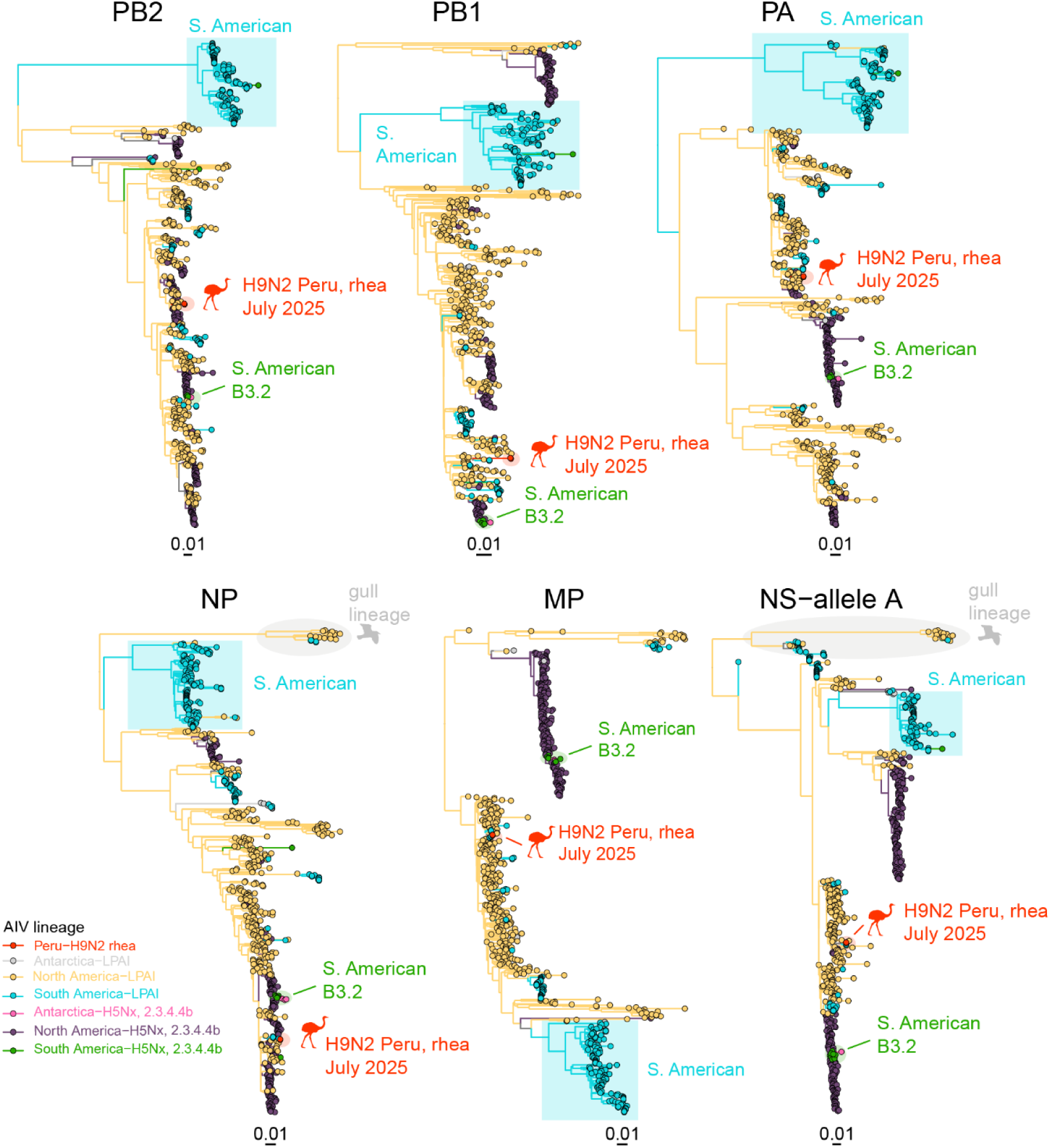
Maximum likelihood subsampled phylogenies of influenza A internal gene segments, including LPAI and HPAI clade 2.3.4.4b viruses collected in the Western hemisphere (e.g., Antarctica, North America and South America). PB2 (*n =* 44,184), PB1 (*n =* 43,802), PA (*n =*44,357), NP (*n =* 44,611), MP (*n =*44,567), and NS-allele A (*n =* 42,238). Tips and branches are colored by AIV lineage: South American LPAI in blue, South American B3.2 in green, Peruvian lesser rhea in red, and gull lineage viruses in segment NS and NP in gray.

**Figure 4.**
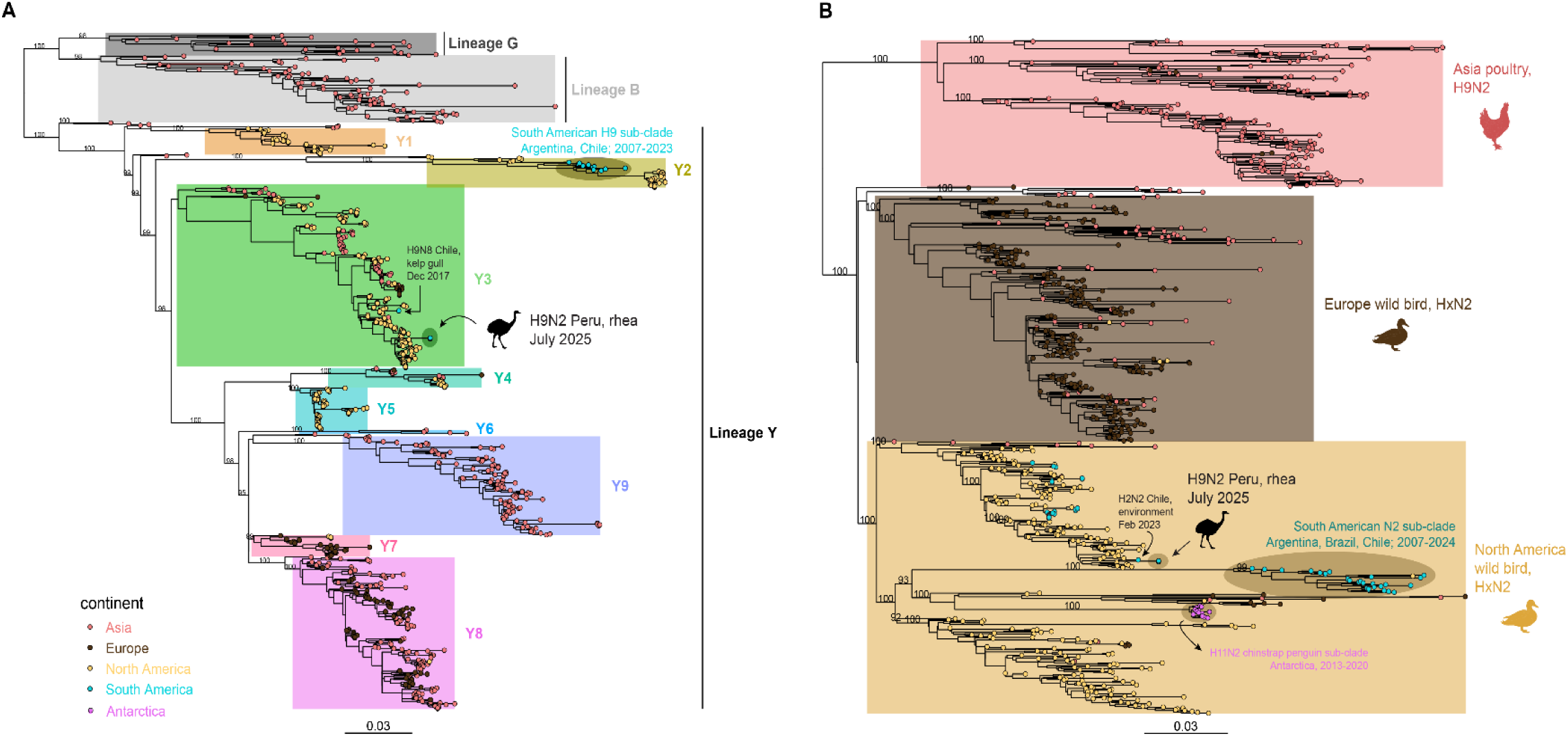
Maximum likelihood phylogenies of HA and NA with samples collected in Asia, Europe, North America and South America. (A) H9Nx ML tree with samples collected from 1974-2025 (*n*= 818). Tips are colored according to the continent of collection, and the phylogeny is annotated with the proposed lineages in Fusaro et al. [24]. Subclades within lineage Y are highlighted as well as two samples (A/green-winged_teal/Wisconsin/228/1976 and A/mallard/Wisconsin/24/1974) classified as Y3 by pairwise distance but clustering with Y2 viruses. UFBoot support values above 90% are displayed on branches leading to key phylogenetic nodes. B. HxN2 ML tree with samples collected from 1961-2025 (*n*= 791). Tips are colored as in the HA tree and well-supported clades (i.e., UFBoot support >90%) are highlighted. UFBoot support values above 90% are displayed on branches leading to key phylogenetic nodes.

### H9N2 rhea viruses represent a new introduction of LPAI-H9N2 into South America

To further elucidate the origin and closest phylogenic relationships of the Peruvian rhea viruses, we took a close look at the HA and NA trees. Analysis of the H9 global phylogeny revealed that the Peruvian rhea viruses do not cluster with other H9 viruses sampled in South America between 2007 and 2023 (Figure 4A, Table 2). Specifically, H9N2 and H9N7 viruses collected from waterfowl, shorebirds and gulls in Chile and Argentina (in blue) cluster together in a “South American H9 sub-clade” within clade 2 of lineage Y (Y2) [24]. Instead, the Peruvian H9N2 rhea viruses position within clade 3 of lineage Y (Y3). Clade Y3 has circulated in North America since at least 1978 in both domestic and wild birds, and it includes 9 different Influenza A subtypes, constituting the largest sub-typic diversity of any defined H9 clade (Table 3). A long branch length separates the Peruvian H9N2 rhea viruses from the most closely related H9 ancestral viruses from North America, suggesting a large gap in AIV surveillance and testing. In an adjacent area of the HA tree, an H9N8 virus collected from a kelp gull in Chile in 2017 positions as a singleton in a different section of clade Y3, confirming the rhea viruses represent an independent (third) introduction of LPAI-H9 from North America into South America. We also looked at the HA cleavage site, which consists of the amino acid sequence PAASDR. This sequence is similar to other LPAI cleavage sites and consistent with its proximity to LPAI strains in the HA tree. Analysis of the N2 global phylogeny shows a similar pattern to the HA tree, with rhea viruses clustering with North American N2 strains rather than within the South American N2 sub-clade (in blue) that includes 28 independent viruses sampled in Argentina, Brazil and Chile between 2007 and 2024 (Figure 4B, Supplemental Table 3). North American N2 viruses have entered South America more frequently than H9, and there is evidence to support an additional introduction into South America, as the rhea viruses cluster with an environmental H2N2 sample collected in Chile in February 2023 (A/environment/Chile/C63085/2023). However, we again see a long branch length separating the rhea viruses from the environmental sample from Chile, confirming a large gap in AIV surveillance and testing. Taken together, the HA and NA global phylogenies, along with the internal gene trees and the sequence of the HA cleavage site, provide support for a new introduction of LPAI-H9N2 of North American origin into South America.

**Table 2.**
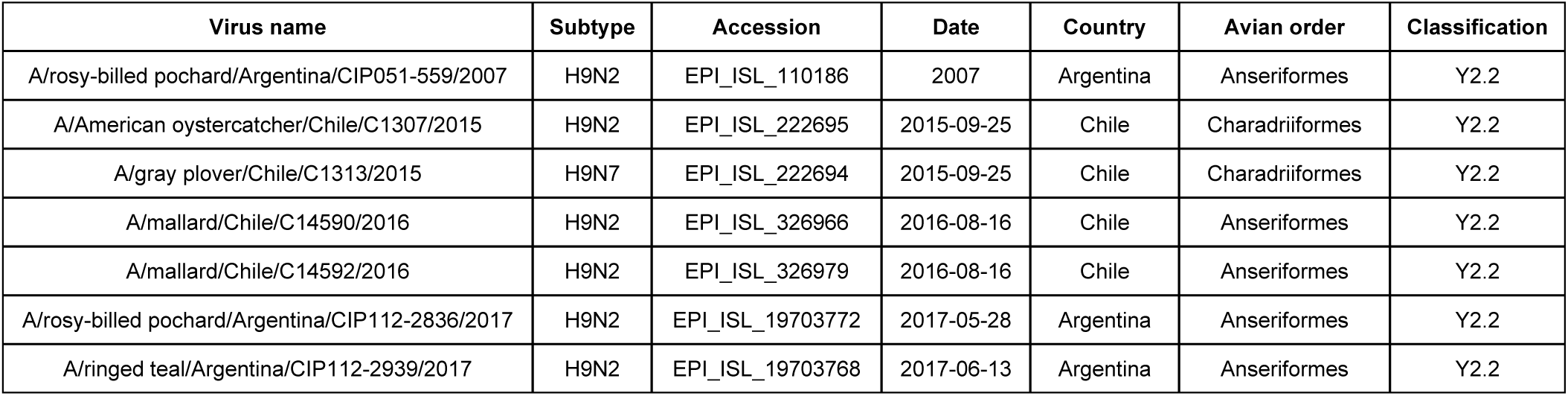
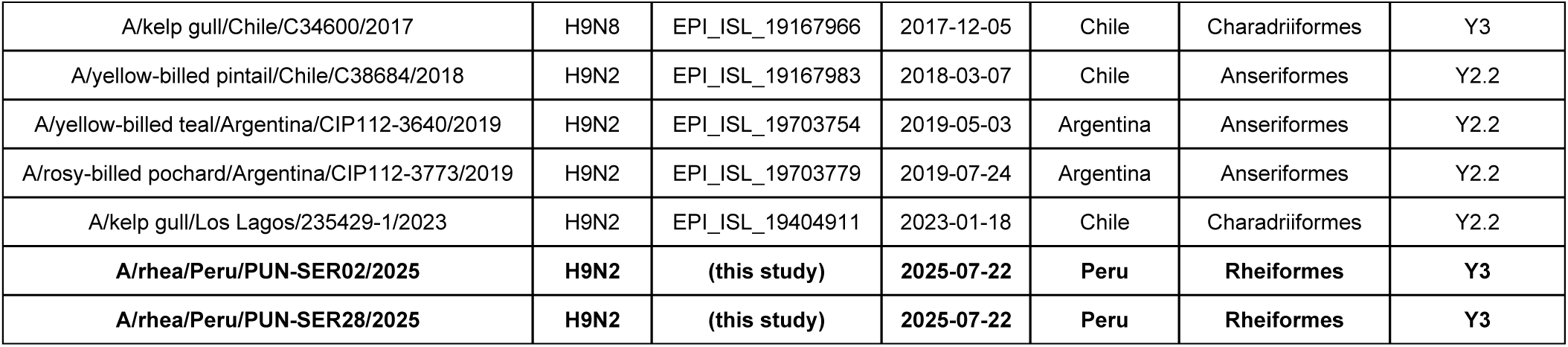
H9Nx viruses sampled in South America, 2007-2025.

| Virus name | Subtype | Accession | Date | Country | Avian order | Classification |
| --- | --- | --- | --- | --- | --- | --- |
| A/rosy-billed pochard/Argentina/CIP051-559/2007 | H9N2 | EPI_ISL_110186 | 2007 | Argentina | Anseriformes | Y2.2 |
| A/American oystercatcher/Chile/C1307/2015 | H9N2 | EPI_ISL_222695 | 2015-09-25 | Chile | Charadriiformes | Y2.2 |
| A/gray plover/Chile/C1313/2015 | H9N7 | EPI_ISL_222694 | 2015-09-25 | Chile | Charadriiformes | Y2.2 |
| A/mallard/Chile/C14590/2016 | H9N2 | EPI_ISL_326966 | 2016-08-16 | Chile | Anseriformes | Y2.2 |
| A/mallard/Chile/C14592/2016 | H9N2 | EPI_ISL_326979 | 2016-08-16 | Chile | Anseriformes | Y2.2 |
| A/rosy-billed pochard/Argentina/CIP112-2836/2017 | H9N2 | EPI_ISL_19703772 | 2017-05-28 | Argentina | Anseriformes | Y2.2 |
| A/ringed teal/Argentina/CIP112-2939/2017 | H9N2 | EPI_ISL_19703768 | 2017-06-13 | Argentina | Anseriformes | Y2.2 |
| A/kelp gull/Chile/C34600/2017 | H9N8 | EPI_ISL_19167966 | 2017-12-05 | Chile | Charadriiformes | Y3 |
| A/yellow-billed pintail/Chile/C38684/2018 | H9N2 | EPI_ISL_19167983 | 2018-03-07 | Chile | Anseriformes | Y2.2 |
| A/yellow-billed teal/Argentina/CIP112-3640/2019 | H9N2 | EPI_ISL_19703754 | 2019-05-03 | Argentina | Anseriformes | Y2.2 |
| A/rosy-billed pochard/Argentina/CIP112-3773/2019 | H9N2 | EPI_ISL_19703779 | 2019-07-24 | Argentina | Anseriformes | Y2.2 |
| A/kelp gull/Los Lagos/235429-1/2023 | H9N2 | EPI_ISL_19404911 | 2023-01-18 | Chile | Charadriiformes | Y2.2 |
| **A/rhea/Peru/PUN-SER02/2025** | **H9N2** | **(this study)** | **2025-07-22** | **Peru** | **Rheiformes** | **Y3** |
| **A/rhea/Peru/PUN-SER28/2025** | **H9N2** | **(this study)** | **2025-07-22** | **Peru** | **Rheiformes** | **Y3** |

**Table 3.** Clade characteristics for H9Nx lineage Y samples including the Peru viruses identified in this study. *Includes 2 samples (A/green-winged_teal/Wisconsin/228/1976 and A/mallard/Wisconsin/24/1974) that are classified under clade Y3 but cluster with clade Y2.

| Clade | Subtypes | Time period | Continent | Country | Host type | # of taxa |
| --- | --- | --- | --- | --- | --- | --- |
| Y1 | H9N1, H9N2, H9N7, H9N9 | 2000-2013 | North America | USA, Mexico | wild avian | 30 |
| Y2 | H9N1, H9N2, H9N6, H9N7, H9N10 | 1993-2025 | North America, South America | Argentina, Chile, USA | domestic avian, wild avian | 36 |
| **Y3** | H9N1, **H9N2**, H9N3, H9N4, H9N5, H9N6, H9N7, H9N8, H9N9 | 1974-2025 | Asia, Europe, North America, South America | **Peru (this study)**, Canada, Chile, China, Germany, Hong Kong, Hungary, Italy, South Korea, Sweden, USA | domestic avian, wild avian | 220* |
| Y4 | H9N1, H9N3, H9N5, H9N7, H9N9 | 2010-2021 | Asia, Europe, North America | Georgia, Poland, Singapore, UK, USA | domestic avian, wild avian | 24 |
| Y5 | H9N1, H9N2, H9N5, H9N7, H9N8, H9N9 | 2003-2007 | North America | USA | wild avian | 51 |
| Y6 | H9N2, H9N6 | 2009-2013 | Asia | Cambodia, Vietnam | domestic avian | 5 |
| Y7 | H9N2, H9N3, H9N7, H9N8, H9N9 | 1993-2010 | Europe, North America | Finland, Germany, Ireland, Italy, Netherlands, Sweden, UK, USA | domestic avian, wild avian | 21 |
| Y8 | H9N1, H9N2, H9N3, H9N4, H9N7, H9N8, H9N9 | 1999-2025 | Asia, Europe, North America, South America | Austria, Bangladesh, Belgium, China, Czech Republic, Finland, France, Germany, Iran, Italy, Japan, Kazakhstan, South Korea, Malaysia, Netherlands, Norway, Poland, Portugal, Russia, Sweden, Switzerland, Taiwan, Thailand, Ukraine, UK, USA, Vietnam | domestic avian, wild avian | 181 |
| Y9 | H9N2 | 1996-2018 | Asia | Malaysia, South Korea | domestic avian, human, swine, wild avian | 115 |

### Mutation analysis

To understand why the animals presented such severe symptoms, including systemic infections and high mortality rates despite being infected with a low pathogenicity strain, we conducted a mutational analysis using FluMut to identify amino acid changes potentially linked to increased pathogenicity, virulence, polymerase activity, receptor binding to α2-6 sialic acid receptors and overall improved viral fitness. We compared the lesser rhea sequences against all complete H9N2 sequences available on GISAID (n ≥ 9,331, accessed on January 8, 2026) to determine whether there were any mutations that were specifically arising in the region or that were exclusive to the lesser rhea samples. A subset of mammalian H5N1 sequences from South America was also included in the analysis to identify any potential mutations reported as mammalian-adapted HPAI variants that have remained uncharacterized to date but that could be highly pathogenic in avian hosts. In total, 44 amino acid substitutions were identified in the lesser rhea sequences (Table 4). Of these, 32 have been previously described, including mutations like M1:I43M and NS-1:V149A that are linked to increased virulence in avian and mammalian hosts. The remaining 12 mutations fall into two categories: 1) 9 have been previously reported, though they remain uncharacterized, including 4 (PA:M86I, HA2-5:A166S, NA:V62I and NS:E26K) that were detected in mammals during the HPAI-H5N1 outbreak in South America that resulted in mass die-offs of birds and mammals [12]; and 2) the remaining 3 are unique to the lesser rhea sequences when compared to all other H9N2 sequences available in public repositories. These mutations include HA2-5:N135H, NS:T58N and a 3-nucleotide insertion that results in the addition of a threonine at position 3 of the HA signal peptide (METTTTLVAILLMVTASNA), which elongates the signal peptide by one amino acid. This new signal peptide, with four threonines in a row, is unique among the 15,000+ sequences queried.

**Table 4.** Mutation table and published associated phenotypes found in lesser rhea samples. Mutations are numbered according to each reference subtype, except HA which is numbered according to subtype H5 and separated by H1 and H2 subunits. Segments are polymerase basic protein 2 (PB2, segment 1), polymerase basic protein (PB1, segment 2), polymerase acidic (PA, segment 3), hemagglutinin (HA, segment 4), nucleoprotein (NP, segment 5), neuraminidase (NA, segment 6), matrix protein (M1, segment 7), nonstructural protein (NS, segment 8). N/a – not applicable. New mutations that are exclusively found in our two rhea sequences are in **bold**.

| # | Segment | Mutation | Subtype | Associated phenotypes | References |
| --- | --- | --- | --- | --- | --- |
| 1 | PB2 | M64T | H9N2 | Highly conserved mutation in human influenza A H1N1, H2N2, and H3N2 viruses | [25,26] |
| 2 |  | V203I | H9N2 | Mutation found in bat influenza H18N11 in Brazil | [27] |
| 3 |  | K389R | H7N9 | Increased polymerase activity in mammalian cells, Increased replication in mammalian cells | [28,29] |
| 4 |  | N448S | H9N2 | Mouse cell line adaptation after serial in vitro passaging | [25] |
| 5 |  | P453S | H9N2 | Uncharacterized mutation | n/a |
| 6 |  | V598T | H7N9 | Increased replication and polymerase activity in mammalian cells, Increased virulence in mice | [28,29] |
| 7 |  | S715N | H5N1 | Decreased virulence in mice | [29,30] |
| 8 |  | L89V, G309D, T339K, R477G, I495V, K627E, A676T | H5N1 | Increased polymerase activity in mammalian cells, Increased virulence in mice | [29,31] |
| 9 | PB1 | D3V | H5N1 | Increased polymerase activity and replication in in avian cells & mammalian cells | [29,32] |
| 10 |  | D622G | H5N1 | Increased polymerase activity in mice, increased polymerase activity in mice | [29,33] |
| 11 | PB1-f2 | N66S | H5N1 | Enhanced antiviral response, replication, and virulence in mice | [29,34,35] |
| 12 | PA | S37A | H7N9 | Increased polymerase activity in mammalian cells | [29,36] |
| 13 |  | V63L | H9N2 | Mutation found in H1N1 sequences | [37,38] |
| 14 |  | M86I | H5N1 | Uncharacterized mutation found in H5N1, including mammals | [12] |
| 15 |  | N325K | H9N2 | Uncharacterized mutation | n/a |
| 16 |  | N383D | H5N1 | Increased polymerase activity in mammalian & avian cells | [29,39,40] |
| 17 |  | N409S | H7N9 | Increased replication and polymerase activity in mammalian cells | [29,36] |
| 18 |  | I465V | H9N2 | Uncharacterized mutation | n/a |
| 19 |  | I504M | H9N2 | Uncharacterized mutation | n/a |
| 20 | HA | **Signal Peptide Insertion 3T** | H9N2 | **New mutation exclusive to rheas in this study** | n/a |
| 21 |  | HA1-5:T54I | H9N2 | Uncharacterized mutation | n/a |
| 22 | | HA1-5:D94N | H5N1 | Increased pseudovirus binding to $\alpha 2$-6 | [29,41] |
| 23 |  | HA1-5:T134A | H9N2 | Increased viral replication in mice lungs and increased virus thermostability | [42] |
| 24 | | HA1-5:S155N | H5N1 | Increased virus binding to $\alpha 2$-6 | [29,43] |
| 25 | | HA1-5:N193K | H5N1 | Increased virus binding to $\alpha 2$-6 | [29,43] |
| 26 |  | HA1-5:I208T | H9N2 | Increased replication in avian and mammalian cells, increased viral replication in mice lungs | [42] |
| 27 |  | HA2-5:K64E | H7N9 | Decreased HA stability, decreased virulence in mice, Increased pH of fusion | [29,44] |
| 28 |  | **HA2-5:N135H** | H9N2 | **New mutation exclusive to rheas in this study** | n/a |
| 29 |  | HA2:A166S | H5N1 | Uncharacterized mutation found in H5N1, including mammals | [12] |
| 30 | | HA1-5:N193K, HA2-5:R167K | H5N1 | Increased virus binding to $\alpha 2$-6 | [29,45] |
| 31 | NP | A184K | H5N1 | Increased replication in avian cells, increased virulence in chickens, enhanced interferon response | [29,46] |
| 32 | NA | V62I | H5N1 | Uncharacterized mutation found in H5N1, including mammals | [12] |
| 33 |  | I117T | H5N1 | Reduced inhibition and susceptibility to Oseltamivir, Zanamivir | [29,47] |
| 34 |  | Y155H | H1N1 | Highly reduced inhibition to Oseltamivir, Zanamivir | [48] |
| 35 | M1 | N30D | H5N1 | Increased virulence in mice | [29,49] |
| 36 |  | I43M | H5N1 | Increased virulence in chickens, ducks, mice | [29,50] |
| 37 |  | T215A | H5N1 | Increased virulence in mice | [29,49] |
| 38 | NS | E26K | H5N1 | Uncharacterized mutation found in H5N1, including mammals | [12] |
| 39 |  | P42S | H5N1 | Decreased antiviral response & increased virulence in mice | [29,51] |
| 40 |  | **T58N** | H9N2 | **New mutation exclusive to rheas in this study** | n/a |
| 41 |  | K55E, K66E, C138F | H5N1 | Decreased interferon response, enhanced replication in mammalian cells | [29,52] |
| 42 |  | I106M | H5N1 | Viral replication in mammalian cells & increased virulence in mice | [29,53–55] |
| 43 |  | L103F | H1N1 | Increased virulence in mice | [29,53] |
| 44 |  | V149A | H5N1 | Decreased interferon response in chickens & Increased virulence in chickens | [29,52] |

## Discussion

The Rhea Conservation Center located in the Andean highlands of Puno-Peru is as a wildlife sanctuary where lesser rheas live in semi-captive conditions, with close monitoring for conservation and research purposes [21]. The Lake Titicaca Binational Special Project has steadily increased the rhea population through captive breeding efforts since the species was declared endangered [56]. The sudden death of multiple animals in June-July 2025 was worrisome and prompted an immediate investigation to determine the cause of death. A total of 46/92 (50%) rheas (24 juveniles and 22 adults) showed symptoms of disease (greenish diarrhea (100%), hyporexia (24%), dyspnea (76%), nasal discharge (42%), drowsiness (18%) and isolation from the flock (73%)), and 45/46 (94%) died (23 juveniles and 22 adults), with a mean survival time of 8 days for juveniles and 19 days for adults, highlighting the role of age in susceptibility and suggesting the likely presence of at least partially protective immunity in older adults. We confirmed the presence of AIV in two animals sampled on July 22^nd^ and the strain was subtyped as LPAI-H9N2. Even though the H9N2 virus was found to be a low pathogenicity strain, we conclude that AIV infection was the likely cause of death due to: 1) symptoms consistent with respiratory, gastro-intestinal and neurological disease, similar to those caused by other AIVs; 2) systemic inflammation and infection, with viral presence in all tissues tested including brain, similar to patterns observed in HPAI infections [57,58]; and 3) presence of mutations associated with increased viral fitness, including new mutations that may have contributed to increased disease severity.

Phylogenetic analysis confirmed that all eight viral genomic segments belonged to the H9N2 subtype of AIVs, that they were of low pathogenic origin from North America, that there were no recombination events, and that the rhea viruses were distinct from other H9N2 viruses previously identified in the region. The HA tree further showed that the rhea viruses clustered within clade Y3 of H9, and that this was phylogenetically distinct from strains sampled in South America from 2007 to 2023 that cluster together within Y2. The only other virus from South America within Y3 was an H9N8 strain collected from a kelp gull in Chile in 2017, but this sample positioned far away in a completely different branch of clade Y3 (Figure 4), suggesting that the rhea viruses represent a third independent introduction of LPAI-H9N2 into South America. Additionally, a long branch separates the rhea viruses from related ancestral HA sequences, suggesting that the strain may have circulated undetected for an extended period in Peru or in a neighboring country prior to detection in lesser rheas in 2025. Given that AIV surveillance and diagnostic capacity in the region is uneven at best, a long period of undetected circulation is plausible [61,62].

We performed a comparative mutation analysis to identify variable sites that might explain the severity of symptoms, widespread dissemination of the virus and high mortality rates observed. Of the 44 mutations identified in this study, several have been associated with increased overall viral fitness in HPAI H5N1 and H7N9 strains, including PB1:D3V, PA:N383D, HA2-5:K64E, NP:A184K, M1:I43M, and NS-1:V149A (Table 5). We also identified 12 additional mutations that remain uncharacterized as they have not been experimentally validated. In this group we find 9 mutations that have been linked to mammalian adaptation, including PA:M86I, HA2-5:A166S, NA:V62I, and NS:E26K, that were found in sea lions from Peru and a human case from Chile during the HPAI-H5N1 outbreak in 2022-2023 [12]. There are 3 additional mutations that are unique to lesser rhea viruses. The most surprising is an insertion of a threonine residue at the third position of the HA signal peptide (METTTTLVAILLMVTASNA). The new unique sequence is not present in any of the ∼15,000+ H9 sequences analyzed, but, except for the additional threonine insertion, it is identical to other H9N2 signal peptides, like from a mallard sampled in Maine in 2023 (EPI_ISL_19239875). Given the importance of HA in viral entry and host adaptation [63], changes within this area are of particular interest since mutations in the HA signal peptide can affect expression levels [64]. Although mutations in this region of the signal peptide have been reported previously [65], the signal peptide length is highly conserved and sequence variation mostly occurs after the signal peptide [66]. Insertions in this region are rare and warrant further investigation to determine their association with viral fitness.

The severe clinical presentation observed in the lesser rheas may not be explained by viral mutations alone. Disease outcomes in AIV infections result from complex interactions between multiple factors beyond viral evolution [67]. The detection of viral genetic material across diverse organ systems including brain, atypical for LPAIs that commonly cluster in respiratory and intestinal tissues [68], suggests possible involvement of host-specific physiological factors and/or potential pathogen-pathogen interactions that increase disease severity. Co-infections of H9N2 with additional viral or bacterial agents have been documented to alter virulence profiles, clinical manifestations, replication dynamics, and tissue tropism in poultry [69]. Although the rhea samples tested negative for Infectious Bronchitis Virus and Avian Paramyxovirus Type 1, it is possible that other undetected viral or bacterial pathogens could have contributed to the severe clinical phenotype observed. Our approach following the initial PCR-based diagnosis of influenza A was to use targeted NGS technology to specifically amplify influenza sequences to quickly characterize the outbreak strain. However, by its nature, this approach limited our ability to detect other pathogens potentially present.

The most likely transmission route and source of infection for this outbreak is through wild birds, especially given that the outbreak location was isolated, remote, at very high altitude (above 4,000 m.a.s.l.) and far away from any poultry operations. Both migratory and resident wild species play important roles in the epidemiology of AIVs [70], and migratory pathways along the Pacific Flyway have been previously linked to the introduction and spread of AIVs from North America into Peru, including during the HPAI-H5N1 outbreak that started in 2022 [12]. *Rhea pennata* shares habitat with *Plegadis ridgwayi* (puna ibis/yanavico), *Charadius alticola* (Puna plover), *Muscisaxicola capistratus* (Cinnamon-bellied Ground-Tyrant), *Anthus furcatus* (Short-billed Pipit) and *Phrygilus alaudinus* (Band-tailed Sierra-Finch). These wild species migrate altitudinally between the highlands above 4,000 m.a.s.l. and the Peruvian coast and are suspected to act as local amplifiers of novel LPAI strains that influence AIV ecology [70,71]. Local wild birds also share more extensive habitats with wild birds that engage in long migrations between North and South America along the Pacific Flyway, providing multiple opportunities for the introduction of AIVs into domestic poultry operations primarily located along the coast [59]. This risk is particularly critical in Peru, as the country has the highest per-capita chicken consumption of Latin America and ranks among the top four countries globally [72], with approximately 55–56 kg consumed per person annually [73]. The poultry sector therefore represents an important component of the national food infrastructure system, and the introduction of pathogenic strains could have considerable economic and public health consequences. During the 2022 HPAI-H5N1 outbreak, more than 37,000 domestic poultry were culled to prevent further spread [74]. Monitoring of seasonal AIV in poultry is a country priority, but testing is limited to H5 and H7 strains. In the case of wildlife and migratory birds, testing remains limited [75].

A larger concern is the potential for H9N2 AIVs to create reassortants with locally circulating strains that could make them especially well adapted to mammals [76]. H9N2 has been reported in bats during routine surveillance efforts in Egypt and South Africa [77,78], further highlighting the host range of this subtype. The introduction of a new strain of H9N2 is therefore of particular concern, as it is a well-recognized donor of internal gene segments that have contributed to the emergence of other influenza strains through reassortment [79]. The limited availability of H9N2 sequences from South America remains a significant challenge for interpreting regional viral evolution. This is the first reported outbreak of LPAI H9N2 in lesser rheas and provides genomic evidence of a distinct introduction of this subtype of AIV into South America. Our findings expand our current knowledge of H9N2 host range in a high altitude environment and provide evidence that low pathogenicity strains can result in high mortality rates, perhaps linked to specific viral mutations. Surveillance programs need to be strengthened to incorporate broad monitoring for circulating AIVs in both poultry and wild birds to enable early detection and close monitoring of regional virus circulation, cross-species transmission, viral evolution, genetic adaptation and future risk assessment

## Materials and Methods

### Sample collection and testing

Samples were collected by trained veterinarians from the Peruvian Wildlife and Forestry Service (SERFOR for its Spanish acronym), the Peruvian Animal Health Service (SENASA), and the Rhea Conservation Center (PEBLT) using standard protocols for biosafety and biocontainment during the performance of their regular duties [80]. One severely clinically ill animal was humanely euthanized to prevent further suffering [81]. Swab samples and tissues were collected into DNA/RNA Shield (Zymo R1200 125) and/or viral transport media (VTM). Nucleic acids were extracted using Quick-DNA/RNA Viral Extraction Kits (Zymo D7021) and tested for influenza A by RT-qPCR using published protocols [82] on a Bio-Rad CFX96 instrument.

### Influenza A subtyping

PCR-positive samples were subtyped using a combination of directed amplification with universal primers targeting conserved genomic regions [83–85], followed by next-generation sequencing. Briefly, RNA samples were reverse transcribed using Superscript IV (Invitrogen 18090050) and amplified using Q5 High-fidelity DNA polymerase (NEB M0491L). Amplification products were prepared into barcoded sequencing libraries using DNA Prep Kits (Illumina 20060059) and barcodes (Illumina 20027213) according to the manufacturer’s instructions. The resulting libraries were quality controlled using High Sensitivity DNA kits (Agilent 5067-4626) on a Bioanalyzer 2100 instrument. Libraries were normalized to 4nM each, pooled, re-quantified using Qubit 1x dsDNA HS Kits (Invitrogen Q33230), and sequenced using Mid Output Sequencing Kits (Illumina FC-420-1004) on an Illumina MiniSeq instrument.

### Bioinformatics

Illumina paired-end raw reads were pre-processed to trim sequencing adaptors and filter out low quality/low complexity reads (Phred scores <Q20, 35 bp minimum length) using Geneious Prime 2025.0.1 and BBDuk [86]. Pre-processed reads were assembled by reference mapping to various HA (H1, H2, H3, H5, H7, H9) and NA (N1, N2, N3, N5, N7, N9) sequences (Accession Numbers: NC_026433, NC_007374, NC_007366, OQ550474, NC_026425, PV985113, OQ550476, PV925107, OP806485, MF046172, OP723829, PV572939) to select influenza-only reads. Filtered reads were then re-assembled *de-novo* using SPAdes [87] to generate complete genomes, and to further confirm subtyping by BLAST. All sequences have been deposited in GenBank Accession # PX488766-PX488781.

### Dataset Curation

To place the Peruvian lesser rhea gene sequences in phylogenetic context, we compiled Influenza A segment datasets from samples collected worldwide by querying the Global Initiative on Sharing All Influenza Data (GISAID; https://gisaid.org/) database [88] (accessed on October 11, 2025). For the H9 and N2 segment datasets, we downloaded all available sequences collected up to October 1, 2025. The resulting datasets consisted of: PB2 (*n=*44,184), PB1 (*n=*43,802), PA (*n=*44,357), H9 (*n=*18,092), NP (*n=* 4,611), N2 (*n=*1,358), MP (*n=*44,567), and NS-allele A (*n=*42,238). Experimental strains were excluded for both segments. Additionally, all human, swine, equine, and canine samples were also excluded from the N2 dataset to remove divergent H1N2 and H3N2 non-avian lineages. For the internal gene segments, only influenza A samples collected between January 1, 2015 and October 1, 2025 with available genomic sequences for all eight gene segments classified as either LPAI or as HPAI H5Nx clade.2.3.4.4b were used. To remove low quality data we filtered out sequences with >5% unresolved bases or <75% of the coding region. Each dataset was aligned using MAFFT [89,90] and trimmed to the coding region. We removed highly divergent sequences that aligned poorly or that introduced frameshifts. Due to small deletions that were difficult to align in the N2 dataset, only sequences ≥1370 bp were included. Divergent NS allele B sequences were also removed.

### Phylogenetic and mutation analysis

Given the large size of the datasets, an initial maximum likelihood (ML) phylogeny was inferred for each segment using FastTree v2.2.0 [91] using a general-time reversible (GTR) with categorical mixture (+CAT) substitution model that accounts for site heterogeneity. To increase the readability of the large phylogenies, ML phylogenetic analyses were also performed on representative, subsampled datasets. The large FastTree ML phylogenies (except for H9) were subsampled using PARNAS, which optimally selects representative taxa to maximize phylogenetic diversity and preserve overall tree topology. The N2 phylogeny was subsampled by constraining the representative taxa selection to 250 sequences from Asia, 250 from Europe, and 250 from North America, while preserving all samples from South America and Antarctica. Since an initial examination of the large FastTree phylogenies showed that the Peruvian rhea samples were not closely related to any Eurasian virus for any of the internal gene segments, these phylogenies were subsampled by selecting a total of 500 representatives that included: 1) viruses from North America (LPAI viruses or HPAI clade 2.3.4.4b viruses); 2) HPAI clade 2.3.4.4b from South America and Antarctica; and 3) all LPAI sequences from South America. Since the H9 phylogeny is largely comprised of Asian sequences, the H9 dataset was subsampled by selecting a random subset of 100 sequences from Asia and keeping all others. To ensure that the closest relatives to each rhea virus segment were preserved in the analysis, a Basic Local Alignment Search Tool (BLAST) search of the GISAID database was performed using the rhea virus segment sequences as queries. The resulting, subsampled datasets consisted of: PB2 (*n=* 685), PB1 (*n=* 673), PA (*n =* 687), H9 (*n =* 818), NP (*n =* 688), N2 (*n =* 719), MP (*n =*694), and NS-allele A (*n =* 642). ML phylogenies were then inferred for each subsampled dataset using IQ-TREE v2.47 [92] with a GTR model of nucleotide substitution with among-site rate heterogeneity modelled through a discretized gamma distribution (+G) and 1,000 ultra-fast bootstrap (UFBOOT) replicates, using the high-performance computational capabilities of the Biowulf Linux cluster at the National Institutes of Health (http://biowulf.nih.gov). Finally, mutation analysis looking for previously characterized amino acid changes related to changes in replication, virulence, transmission, antiviral resistance and other phenotypes was done using FluMutGUI v3.2.0 [93], and mutation frequency was determined by comparison among all H9N2 gene segments available to date on GISAID (accessed on January 8, 2026), which included over 9,330 H9N2 sequences.

## Acknowledgments

The authors thank Alberto Mamani and Elmer Ventura from ATFFS-Puno for support during field work, and Bailón Sacachipana from SENASA-Puno for support with sample transport from field to lab. We also acknowledge SENASA officials for their informal contribution to the discussion of the findings. Finally, we gratefully acknowledge all data contributions (including authors and originating labs responsible for obtaining specimens and generating sequences) to the GISAID EpiFlu database from which reference sequences were derived for this research

## References

1. Peacock TP, James J, Sealy JE, Iqbal M. A Global Perspective on H9N2 Avian Influenza Virus. Viruses. 2019;11:620. 10.3390/v11070620

2. Jallow MM, Barry MA, Fall A, Ndiaye NK, Kiori D, Sy S, et al. Influenza A Virus in Pigs in Senegal and Risk Assessment of Avian Influenza Virus (AIV) Emergence and Transmission to Human. Microorganisms. 2023;11:1961. 10.3390/microorganisms11081961

3. Yong-feng Z, Fei-fei D, Jia-yu Y, Feng-xia Z, Chang-qing J, Jian-li W, et al. Intraspecies and interspecies transmission of mink H9N2 influenza virus. Sci Rep. 2017;7:7429. 10.1038/s41598-017-07879-1

4. Yang J, Yan J, Zhang C, Li S, Yuan M, Zhang C, et al. Genetic, biological and epidemiological study on a cluster of H9N2 avian influenza virus infections among chickens, a pet cat, and humans at a backyard farm in Guangxi, China. Emerg Microbes Infect. 2023;12. 10.1080/22221751.2022.2143282

5. Liu K, Wang X, Sun Y, Liu X, Wang X. Will avian influenza virus (H10N3) cause a major public health threat? Animals and Zoonoses. Elsevier BV; 2025; 10.1016/j.azn.2025.11.003

6. Pu J, Wang S, Yin Y, Zhang G, Carter RA, Wang J, et al. Evolution of the H9N2 influenza genotype that facilitated the genesis of the novel H7N9 virus. Proc Natl Acad Sci U S A. National Academy of Sciences; 2015;112:548–53. 10.1073/pnas.1422456112

7. Liu J, Zhang J, Huang F, Zhang Y, Luo H, Zhang H. Complex reassortment of polymerase genes in Asian influenza A virus H7 and H9 subtypes. Infection, Genetics and Evolution. 2014;23:203–8. 10.1016/j.meegid.2014.02.016

8. Guan YI, Shortridge KF, Krauss S, Webster RG. Molecular characterization of H9N2 influenza viruses: Were they the donors of the “‘internal’” genes of H5N1 viruses in Hong Kong? 1999. 10.1073/pnas.96.16.9363

9. World Health Organization. WHO Avian Influenza Weekly Update # 1029 [Internet]. 2026 Jan. https://cdn.who.int/media/docs/default-source/wpro---documents/emergency/surveillance/avian-influenza/ai_20260116.pdf?sfvrsn=125a11a2_1&download=true. Accessed 18 Jan 2026

10. Senne DA. Avian Influenza in North and South America, 2002–2005. Avian Dis. 2007;51:167–73. 10.1637/7621-042606R1.1

11. Harvey JA, Mullinax JM, Runge MC, Prosser DJ. The changing dynamics of highly pathogenic avian influenza H5N1: Next steps for management & science in North America. Biol Conserv. 2023;282:110041. 10.1016/j.biocon.2023.110041

12. Leguia M, Garcia-Glaessner A, Muñoz-Saavedra B, Juarez D, Barrera P, Calvo-Mac C, et al. Highly pathogenic avian influenza A (H5N1) in marine mammals and seabirds in Peru. Nat Commun. 2023;14:5489. 10.1038/s41467-023-41182-0

13. Krammer F, Hermann E, Rasmussen AL. Highly pathogenic avian influenza H5N1: history, current situation, and outlook. J Virol. 2025;99:e0220924. 10.1128/jvi.02209-24

14. Vanstreels RET, Nelson MI, Artuso MC, Marchione VD, Piccini LE, Benedetti E, et al. Novel Highly Pathogenic Avian Influenza A(H5N1) Virus, Argentina, 2025. Emerg Infect Dis [Internet]. 2025;31. 10.3201/eid3112.250783

15. Sevilla N, Lizarraga W, Jimenez-Vasquez V, Hurtado V, Molina IS, Huarca L, et al. Highly pathogenic avian influenza A (H5N1) virus outbreak in Peru in 2022–2023. Infectious Medicine. 2024;3:100108. 10.1016/j.imj.2024.100108

16. Nelson MI, Pollett S, Ghersi B, Silva M, Simons MP, Icochea E, et al. The genetic diversity of influenza A viruses in wild birds in Peru. PLoS One. Public Library of Science; 2016;11. 10.1371/journal.pone.0146059

17. Servicio Nacional Forestal y de Fauna Silvestre (SERFOR). Libro Rojo de la Fauna Silvestre Amenazada del Perú. 2018 [cited 2026 Jan 18]; https://cdn.www.gob.pe/uploads/document/file/1269071/Libro-Rojo.pdf?v=1598652288. Accessed 18 Jan 2026

18. Diario Oficial El Peruano. Decreto Supremo N.° 034-2004-AG - Categorización de Especies Amenazadas de Fauna Silvestre [Internet]. Lima; 2004. https://www.gob.pe/institucion/osinfor/normas-legales/792194-034-2004-ag-categorizacion-de-especies-amenazadas-de-fauna-silvestre

19. Ministerio de Agricultura y Riego. Situación poblacional del Suri en el Perú: Resultados del II Censo Nacional. 2018 Dec.

20. Proyecto Especial Binacional Lago Titicaca (PEBLT). Conservación del Suri (Rhea pennata) avances y logros [Internet]. 2017 [cited 2026 Jan 18]. https://www.gob.pe/institucion/peblt/informes-publicaciones/459750-conservacion-del-suri-rhea-penata-avances-y-logros-1ra-edicion-peblt. Accessed 18 Jan 2026

21. Proyecto Especial Binacional Lago Titicaca – PELT. Avances en el manejo y la conservacion del suri [Internet]. 2008. https://sinia.minam.gob.pe/documentos/avances-manejo-conservacion-suri. Accessed 19 Jan 2026

22. Panigrahy B, Senne DA, Pearson JE. Presence of Avian Influenza Virus (AIV) Subtypes H5N2 and H7N1 in Emus (Dromaius novaehollandiae) and Rheas (Rhea americana): Virus Isolation and Serologic Findings. Avian Dis. 1995;39:64. 10.2307/1591983

23. Coombes HA, Terrey J, Schlachter A-L, McCarter P, Regina I, Hepple R, et al. Infection of ratites with clade 2.3.4.4b HPAIV H5N1: Potential implications for zoonotic risk. 2025. 10.1101/2025.09.08.674895

24. Fusaro A, Pu J, Zhou Y, Lu L, Tassoni L, Lan Y, et al. Proposal for a Global Classification and Nomenclature System for A/H9 Influenza Viruses. Emerg Infect Dis. 2024;30. 10.3201/eid3008.231176

25. Liang L, Jiang L, Li J, Zhao Q, Wang J, He X, et al. Low Polymerase Activity Attributed to PA Drives the Acquisition of the PB2 E627K Mutation of H7N9 Avian Influenza Virus in Mammals. mBio. 2019;10. 10.1128/mBio.01162-19

26. Wen L, Chu H, Wong BH-Y, Wang D, Li C, Zhao X, et al. Large-scale sequence analysis reveals novel human-adaptive markers in PB2 segment of seasonal influenza A viruses. Emerg Microbes Infect. 2018;7:1–12. 10.1038/s41426-018-0050-0

27. Campos ACA, Góes LGB, Moreira-Soto A, de Carvalho C, Ambar G, Sander A-L, et al. Bat Influenza A(HL18NL11) Virus in Fruit Bats, Brazil. Emerg Infect Dis. 2019;25:333–7. 10.3201/eid2502.181246

28. Hu M, Yuan S, Zhang K, Singh K, Ma Q, Zhou J, et al. PB2 substitutions V598T/I increase the virulence of H7N9 influenza A virus in mammals. Virology. 2017;501:92–101. 10.1016/j.virol.2016.11.008

29. Suttie A, Deng Y-M, Greenhill AR, Dussart P, Horwood PF, Karlsson EA. Inventory of molecular markers affecting biological characteristics of avian influenza A viruses. Virus Genes. 2019;55:739–68. 10.1007/s11262-019-01700-z

30. Sun H, Cui P, Song Y, Qi Y, Li X, Qi W, et al. PB2 segment promotes high-pathogenicity of H5N1 avian influenza viruses in mice. Front Microbiol. 2015;6. 10.3389/fmicb.2015.00073

31. Li J, Ishaq M, Prudence M, Xi X, Hu T, Liu Q, et al. Single mutation at the amino acid position 627 of PB2 that leads to increased virulence of an H5N1 avian influenza virus during adaptation in mice can be compensated by multiple mutations at other sites of PB2. Virus Res. 2009;144:123–9. 10.1016/j.virusres.2009.04.008

32. Elgendy EM, Arai Y, Kawashita N, Daidoji T, Takagi T, Ibrahim MS, et al. Identification of polymerase gene mutations that affect viral replication in H5N1 influenza viruses isolated from pigeons. Journal of General Virology. 2017;98:6–17. 10.1099/jgv.0.000674

33. Feng X, Wang Z, Shi J, Deng G, Kong H, Tao S, et al. Glycine at Position 622 in PB1 Contributes to the Virulence of H5N1 Avian Influenza Virus in Mice. J Virol. 2016;90:1872–9. 10.1128/JVI.02387-15

34. Conenello GM, Zamarin D, Perrone LA, Tumpey T, Palese P. A Single Mutation in the PB1-F2 of H5N1 (HK/97) and 1918 Influenza A Viruses Contributes to Increased Virulence. PLoS Pathog. 2007;3:e141. 10.1371/journal.ppat.0030141

35. Schmolke M, Manicassamy B, Pena L, Sutton T, Hai R, Varga ZT, et al. Differential Contribution of PB1-F2 to the Virulence of Highly Pathogenic H5N1 Influenza A Virus in Mammalian and Avian Species. PLoS Pathog. 2011;7:e1002186. 10.1371/journal.ppat.1002186

36. Yamayoshi S, Yamada S, Fukuyama S, Murakami S, Zhao D, Uraki R, et al. Virulence-Affecting Amino Acid Changes in the PA Protein of H7N9 Influenza A Viruses. J Virol. 2014;88:3127–34. 10.1128/JVI.03155-13

37. Badar N, Ikram A, Salman M, Saeed S, Mirza HA, Ahad A, et al. Evolutionary analysis of seasonal influenza A viruses in Pakistan 2020–2023. Influenza Other Respir Viruses. 2024;18. 10.1111/irv.13262

38. Swanson NJ, Marinho P, Dziedzic A, Jedlicka A, Liu H, Fenstermacher K, et al. 2019–2020 H1N1 clade A5a.1 viruses have better in vitro fitness compared with the co-circulating A5a.2 clade. Sci Rep. 2023;13:10223. 10.1038/s41598-023-37122-z

39. Song J, Xu J, Shi J, Li Y, Chen H. Synergistic Effect of S224P and N383D Substitutions in the PA of H5N1 Avian Influenza Virus Contributes to Mammalian Adaptation. Sci Rep. 2015;5:10510. 10.1038/srep10510

40. Song J, Feng H, Xu J, Zhao D, Shi J, Li Y, et al. The PA Protein Directly Contributes to the Virulence of H5N1 Avian Influenza Viruses in Domestic Ducks. J Virol. 2011;85:2180–8. 10.1128/JVI.01975-10

41. Su Y, Yang H-Y, Zhang B-J, Jia H-L, Tien P. Analysis of a point mutation in H5N1 avian influenza virus hemagglutinin in relation to virus entry into live mammalian cells. Arch Virol. 2008;153:2253–61. 10.1007/s00705-008-0255-y

42. Zhang J, Wang X, Chen Y, Ye H, Ding S, Zhang T, et al. Mutational antigenic landscape of prevailing H9N2 influenza virus hemagglutinin spectrum. Cell Rep. Elsevier B.V.; 2023;42. 10.1016/j.celrep.2023.113409

43. Wang W, Lu B, Zhou H, Suguitan AL, Cheng X, Subbarao K, et al. Glycosylation at 158N of the Hemagglutinin Protein and Receptor Binding Specificity Synergistically Affect the Antigenicity and Immunogenicity of a Live Attenuated H5N1 A/Vietnam/1203/2004 Vaccine Virus in Ferrets. J Virol. 2010;84:6570–7. 10.1128/JVI.00221-10

44. Sun X, Belser JA, Yang H, Pulit-Penaloza JA, Pappas C, Brock N, et al. Identification of key hemagglutinin residues responsible for cleavage, acid stability, and virulence of fifth-wave highly pathogenic avian influenza A(H7N9) viruses. Virology. 2019;535:232–40. 10.1016/j.virol.2019.07.012

45. Yamada S, Suzuki Y, Suzuki T, Le MQ, Nidom CA, Sakai-Tagawa Y, et al. Haemagglutinin mutations responsible for the binding of H5N1 influenza A viruses to human-type receptors. Nature. 2006;444:378–82. 10.1038/nature05264

46. Wasilenko JL, Sarmento L, Pantin-Jackwood MJ. A single substitution in amino acid 184 of the NP protein alters the replication and pathogenicity of H5N1 avian influenza viruses in chickens. Arch Virol. 2009;154:969–79. 10.1007/s00705-009-0399-4

47. Kode SS, Pawar SD, Tare DS, Keng SS, Hurt AC, Mullick J. A novel I117T substitution in neuraminidase of highly pathogenic avian influenza H5N1 virus conferring reduced susceptibility to oseltamivir and zanamivir. Vet Microbiol. 2019;235:21–4. 10.1016/j.vetmic.2019.06.005

48. Monto AS, McKimm-Breschkin JL, Macken C, Hampson AW, Hay A, Klimov A, et al. Detection of Influenza Viruses Resistant to Neuraminidase Inhibitors in Global Surveillance during the First 3 Years of Their Use. Antimicrob Agents Chemother. 2006;50:2395–402. 10.1128/AAC.01339-05

49. Fan S, Deng G, Song J, Tian G, Suo Y, Jiang Y, et al. Two amino acid residues in the matrix protein M1 contribute to the virulence difference of H5N1 avian influenza viruses in mice. Virology. 2009;384:28–32. 10.1016/j.virol.2008.11.044

50. Nao N, Kajihara M, Manzoor R, Maruyama J, Yoshida R, Muramatsu M, et al. A Single Amino Acid in the M1 Protein Responsible for the Different Pathogenic Potentials of H5N1 Highly Pathogenic Avian Influenza Virus Strains. PLoS One. 2015;10:e0137989. 10.1371/journal.pone.0137989

51. Jiao P, Tian G, Li Y, Deng G, Jiang Y, Liu C, et al. A Single-Amino-Acid Substitution in the NS1 Protein Changes the Pathogenicity of H5N1 Avian Influenza Viruses in Mice. J Virol. 2008;82:1146–54. 10.1128/JVI.01698-07

52. Li J, Zhang K, Chen Q, Zhang X, Sun Y, Bi Y, et al. Three amino acid substitutions in the NS1 protein change the virus replication of H5N1 influenza virus in human cells. Virology. 2018;519:64–73. 10.1016/j.virol.2018.04.004

53. Ayllon J, Domingues P, Rajsbaum R, Miorin L, Schmolke M, Hale BG, et al. A Single Amino Acid Substitution in the Novel H7N9 Influenza A Virus NS1 Protein Increases CPSF30 Binding and Virulence. J Virol. 2014;88:12146–51. 10.1128/JVI.01567-14

54. Kuo R-L, Krug RM. Influenza A Virus Polymerase Is an Integral Component of the CPSF30-NS1A Protein Complex in Infected Cells. J Virol. 2009;83:1611–6. 10.1128/JVI.01491-08

55. Spesock A, Malur M, Hossain MJ, Chen L-M, Njaa BL, Davis CT, et al. The Virulence of 1997 H5N1 Influenza Viruses in the Mouse Model Is Increased by Correcting a Defect in Their NS1 Proteins. J Virol. 2011;85:7048–58. 10.1128/JVI.00417-11

56. Servicio Nacional Forestal y de Fauna Silvestre (SERFOR). Plan Nacional pata la Conservacion del Suri (Rhea pennata) Periodo 2015-2020 [Internet]. 2015. https://www.gob.pe/institucion/serfor/informes-publicaciones/1124090-plan-nacional-para-la-conservacion-del-suri-rhea-pennata. Accessed 18 Jan 2026

57. Floyd T, Banyard AC, Lean FZX, Byrne AMP, Fullick E, Whittard E, et al. Encephalitis and Death in Wild Mammals at a Rehabilitation Center after Infection with Highly Pathogenic Avian Influenza A(H5N8) Virus, United Kingdom. Emerg Infect Dis. 2021;27:2856–63. 10.3201/eid2711.211225

58. Korteweg C, Gu J. Pathology, Molecular Biology, and Pathogenesis of Avian Influenza A (H5N1) Infection in Humans. Am J Pathol. 2008;172:1155–70. 10.2353/ajpath.2008.070791

59. Olsen B, Munster VJ, Wallensten A, Waldenström J, Osterhaus ADME, Fouchier RAM. Global Patterns of Influenza A Virus in Wild Birds. Science (1979). 2006;312:384–8. 10.1126/science.1122438

60. Webster RG, Bean WJ, Gorman OT, Chambers TM, Kawaoka Y. Evolution and ecology of influenza A viruses. Microbiol Rev. 1992;56:152–79. 10.1128/mr.56.1.152-179.1992

61. Grijalva-Trejo X, Vasquez C. Highly pathogenic Avian Influenza (H5N1) in Latin America: Epidemiology, control strategies, and One Health implications. Letters In Animal Biology. 2026;09–18. 10.62310/liab.v6i1.268

62. Neumann G, Kawaoka Y. Predicting the Next Influenza Pandemics. J Infect Dis. 2019;219:S14–20. 10.1093/infdis/jiz040

63. Xu R, Wilson IA. Structural characterization of an early fusion intermediate of influenza virus hemagglutinin. J Virol. 2011;85:5172–82. 10.1128/JVI.02430-10

64. Doud MB, Lee JM, Bloom JD. How single mutations affect viral escape from broad and narrow antibodies to H1 influenza hemagglutinin. Nat Commun. 2018;9:1386. 10.1038/s41467-018-03665-3

65. Canale AS, Venev S V., Whitfield TW, Caffrey DR, Marasco WA, Schiffer CA, et al. Synonymous Mutations at the Beginning of the Influenza A Virus Hemagglutinin Gene Impact Experimental Fitness. J Mol Biol. 2018;430:1098–115. 10.1016/j.jmb.2018.02.009

66. Burke DF, Smith DJ. A recommended numbering scheme for influenza A HA subtypes. PLoS One. 2014;9:e112302. 10.1371/journal.pone.0112302

67. Dobson A. The Evolution of Virulence: An Ecological Perspective. Annu Rev Virol. 2025;12:135–56. 10.1146/annurev-virology-093022-020712

68. Böttcher-Friebertshäuser E, Garten W, Matrosovich M, Klenk HD. The Hemagglutinin: A Determinant of Pathogenicity. 2014. p. 3–34. 10.1007/82_2014_384

69. Samy A, Naguib M. Avian Respiratory Coinfection and Impact on Avian Influenza Pathogenicity in Domestic Poultry: Field and Experimental Findings. Vet Sci. 2018;5:23. 10.3390/vetsci5010023

70. Chen LL, Ip JD, Chan WM, Lam SJA, Leung RCY, Yip CCY, et al. Enhanced replication of a contemporary avian influenza A H9N2 virus in human respiratory organoids. Emerg Microbes Infect. Taylor and Francis Ltd.; 2025;14. 10.1080/22221751.2025.2576574

71. Rademan R, Geldenhuys M, Markotter W. Detection and Characterization of an H9N2 Influenza A Virus in the Egyptian Rousette Bat in Limpopo, South Africa. Viruses. MDPI; 2023;15. 10.3390/v15020498

72. Kandeil A, Gomaa MR, Shehata MM, El Taweel AN, Mahmoud SH, Bagato O, et al. Isolation and Characterization of a Distinct Influenza A Virus from Egyptian Bats. 2019; 10.1128/JVI

73. Verhagen JH, van Dijk JGB, Vuong O, Bestebroer T, Lexmond P, Klaassen M, et al. Migratory Birds Reinforce Local Circulation of Avian Influenza Viruses. PLoS One. 2014;9:e112366. 10.1371/journal.pone.0112366

74. Gibbons RE, Benham PM. Notes on birds of the high Andes of Peru. Ornitología Colombiana. 2021;76–86. 10.59517/oc.e248

75. Asociacion Peruana de Avicultura (APA). Boletin APA Informa Edicion 013. https://apa.org.pe/portfolio-item/boletin-setiembre-2024/. 2024.

76. International Trade Administration USD of CITA. Peru Country Commercial Guide. https://www.trade.gov/country-commercial-guides/peru-agriculture-sectors. 2023.

77. Editora Andina. Gripe aviar: sacrifican aves de corral en Huacho y Chiclayo para evitar más contagios. https://andina.pe/ingles/noticia-gripe-aviar-sacrifican-aves-corral-huacho-y-chiclayo-para-evitar-mas-contagios-919928.aspx. 2022.

78. Nelson MI, Pollett S, Ghersi B, Silva M, Simons MP, Icochea E, et al. The Genetic Diversity of Influenza A Viruses in Wild Birds in Peru. PLoS One. 2016;11:e0146059. 10.1371/journal.pone.0146059

79. Cargnin Faccin F, Perez DR. Pandemic preparedness through vaccine development for avian influenza viruses. Hum Vaccin Immunother. 2024;20. 10.1080/21645515.2024.2347019

80. Miller LP, Miknis RA, Flory GA. Carcass management guidelines Effective disposal of animal carcasses and contaminated materials on small to medium-sized farms FAO AnimAl prOduCtiOn And hEAlth / guidelines 23.

81. Leary SL. AVMA guidelines for the euthanasia of animals : 2020 edition. American Veterinary Medical Association; 2020.

82. Shu B, Kirby MK, Davis WG, Warnes C, Liddell J, Liu J, et al. Multiplex real-time reverse transcription PCR for influenza a virus, influenza b virus, and severe acute respiratory syndrome coronavirus 2. Emerg Infect Dis. Centers for Disease Control and Prevention (CDC); 2021;27:1821–30. 10.3201/eid2707.210462

83. Hoffmann E, Stech J, Guan Y, Webster RG, Perez DR. Universal primer set for the full-length amplification of all influenza A viruses. Arch Virol. 2001;146:2275–89. 10.1007/s007050170002

84. Zhou B, Donnelly ME, Scholes DT, St. George K, Hatta M, Kawaoka Y, et al. Single-Reaction Genomic Amplification Accelerates Sequencing and Vaccine Production for Classical and Swine Origin Human Influenza A Viruses. J Virol. American Society for Microbiology; 2009;83:10309–13. 10.1128/jvi.01109-09

85. Hurtado R, Fabrizio T, Vanstreels RTE, Krauss S, Webby RJ, Webster RG, et al. Molecular characterization of subtype H11N9 avian influenza virus isolated from shorebirds in Brazil. PLoS One. Public Library of Science; 2015;10. 10.1371/journal.pone.0145627

86. Bushnell B. BBMap short read aligner, and other bioinformatic tools. https://sourceforge.net/projects/bbmap/. 2022.

87. Bankevich A, Nurk S, Antipov D, Gurevich AA, Dvorkin M, Kulikov AS, et al. SPAdes: A new genome assembly algorithm and its applications to single-cell sequencing. Journal of Computational Biology. 2012;19:455–77. 10.1089/cmb.2012.0021

88. Shu Y, McCauley J. GISAID: Global initiative on sharing all influenza data – from vision to reality. Eurosurveillance. 2017;22. 10.2807/1560-7917.ES.2017.22.13.30494

89. Katoh K, Standley DM. MAFFT Multiple Sequence Alignment Software Version 7: Improvements in Performance and Usability. Mol Biol Evol. 2013;30:772–80. 10.1093/molbev/mst010

90. Katoh K, Toh H. Recent developments in the MAFFT multiple sequence alignment program. Brief Bioinform. 2008;9:286–98. 10.1093/bib/bbn013

91. Price MN, Dehal PS, Arkin AP. FastTree 2 – Approximately Maximum-Likelihood Trees for Large Alignments. PLoS One. 2010;5:e9490. 10.1371/journal.pone.0009490

92. Minh BQ, Schmidt HA, Chernomor O, Schrempf D, Woodhams MD, von Haeseler A, et al. IQ-TREE 2: New Models and Efficient Methods for Phylogenetic Inference in the Genomic Era. Mol Biol Evol. 2020;37:1530–4. 10.1093/molbev/msaa015

93. Giussani E, Sartori A, Salomoni A, Cavicchio L, de Battisti C, Pastori A, et al. FluMut: a tool for mutation surveillance in highly pathogenic H5N1 genomes. Virus Evol. 2025;11. 10.1093/ve/veaf011

